# Evidence of the Recombinant Origin and Ongoing Mutations in Severe Acute Respiratory Syndrome Coronavirus 2 (SARS-CoV-2)

**DOI:** 10.1101/2020.03.16.993816

**Authors:** Jiao-Mei Huang, Syed Sajid Jan, Xiaobin Wei, Yi Wan, Songying Ouyang

**Author notes:** Contributed equally to this work (Jiao-Mei Huang, Syed Sajid Jan, Songying Ouyang). To whom correspondence should be addressed (SO); (YW).

## Abstract

The recent global outbreak of viral pneumonia designated as Coronavirus Disease 2019 (COVID-19) by coronavirus (SARS-CoV-2) has threatened global public health and urged to investigate its source. Whole genome analysis of SARS-CoV-2 revealed ~96% genomic similarity with bat CoV (RaTG13) and clustered together in phylogenetic tree. Furthermore, RaTG13 also showed 97.43% spike protein similarity with SARS-CoV-2 suggesting that RaTG13 is the closest strain. However, RBD and key amino acid residues supposed to be crucial for human-to-human and cross-species transmission are homologues between SARS-CoV-2 and pangolin CoVs. These results from our analysis suggest that SARS-CoV-2 is a recombinant virus of bat and pangolin CoVs. Moreover, this study also reports mutations in coding regions of 125 SARS-CoV-2 genomes signifying its aptitude for evolution. In short, our findings propose that homologous recombination has been occurred between bat and pangolin CoVs that triggered cross-species transmission and emergence of SARS-CoV-2, and, during the ongoing outbreak, SARS-CoV-2 is still evolving for its adaptability.

## INTRODUCTION

The family *Coronaviridae* is comprised of large, enveloped, single stranded, and positive-sense RNA viruses that can infect a wide range of animals including humans (To *et al*., 2013; Guan *et al*., 2003). The viruses are further classified into four genera: *alpha, beta, gamma,* and *delta* coronavirus (King *et al.,* 2012). So far, all coronaviruses (CoVs) identified in human belong to the genera *alpha* and *beta.* Among them betaCoVs are of particular importance. Different novel strains of highly infectious betaCoVs have been emerged in human populations in the past two decades that have caused severe health concern all over the world. Severe acute respiratory syndrome coronavirus (SARS-CoV) was first recognized in 2003, causing a global outbreak (Zhong, 2004; Peiris *et al.,* 2004; Cherry, 2004). It was followed by another pandemic event in 2012 by a novel strain of coronavirus designated as Middle East respiratory syndrome coronavirus (MERS-CoV) (Lu *et al.,* 2013). Both CoVs were zoonotic pathogens and evolved in animals. Bats in the genus *Rhinolophus* are natural reservoir of coronaviruses worldwide, and it is presumed that both SARS-CoV and MERS-CoV have been transmitted to human through some intermediate mammalian hosts (Li *et al.,* 2005a; Bolles *et al.,* 2011; Al-Tawfiq and Memish, 2014). Recently, emergence of another pandemic termed as Coronavirus Disease 2019 (COVID-19) by World Health Organization (WHO) caused by a novel severe acute respiratory syndrome coronavirus 2 (SARS-CoV-2) has been reported (Zhu *et al.,* 2020). To date, more than 174,000 people are infected and over 6,600 death tolls, having transmission clusters worldwide including China, Italy, South Korea, Iran, Japan, USA, France, Spain, Germany and several other countries causing alarming global health concern.

The large trimeric spike glycoprotein (S) located on the surface of CoVs is crucial for viral infection and pathogenesis, which is further subdivided into N-terminal S1 subunit and C-terminal S2 domain. The S1 subunit is specialized in recognizing receptors on host cell, comprising of two separate domains located at N- and C-terminal which can fold independently and facilitate receptor engagement (Masters, 2006). Receptor-binding domains (RBDs) of most CoVs are located on S1 C-terminus and enable attachment to its host receptor (Li *et al.,* 2005b). The host specificity of virus particle is determined by amino acid sequence of RBD and is usually dissimilar among different CoVs. Therefore, RBD is a core determinant for tissue tropism and host range of CoVs. This article presents SARS-CoV-2 phylogenetic trees, comparison and analysis of genome, spike protein, and RBD amino acid sequences of different CoVs, deducing source and etiology of COVID-19 and evolutionary relationship among SARS-CoV-2 in human.

## RESULTS AND DISCUSSION

### Phylogenetic classification of SARS-CoV-2 and its closely related CoVs

To determine the evolutionary relationship of the SARS-CoV-2, phylogenetic analysis was performed on whole genomic sequences of different CoVs from various hosts. The Maximum-likelihood (ML) phylogenetic tree is shown in **Figure 1**, which illustrates four main groups representing four genera of CoVs, *alpha, beta, gamma,* and *delta.* In the phylogenetic tree, strains of SARS-CoV-2 (red colored) are cluster together and belong to the genera Betacoronavirus. Among Beta-CoVs, SARS-CoV, Civet SARS CoV, Bat SARS-like CoVs, bat/RaTG13 CoV, and SARS-CoVs-2 clustered together forming a discrete clade from MERS-CoVs. The clade is further divided into two branches and one of the branches comprises all SARS-CoV-2 strains clustered together with Bat/Yunnan/RaTG13 CoV forming a monophyletic group. Bat/Yunnan/RaTG13 exhibited ~96% genomic similarity with SARS-CoV-2. This specifies that SARS-CoV-2 is closely related to Bat/Yunnan/RaTG13 CoV.

**Figure 1.**
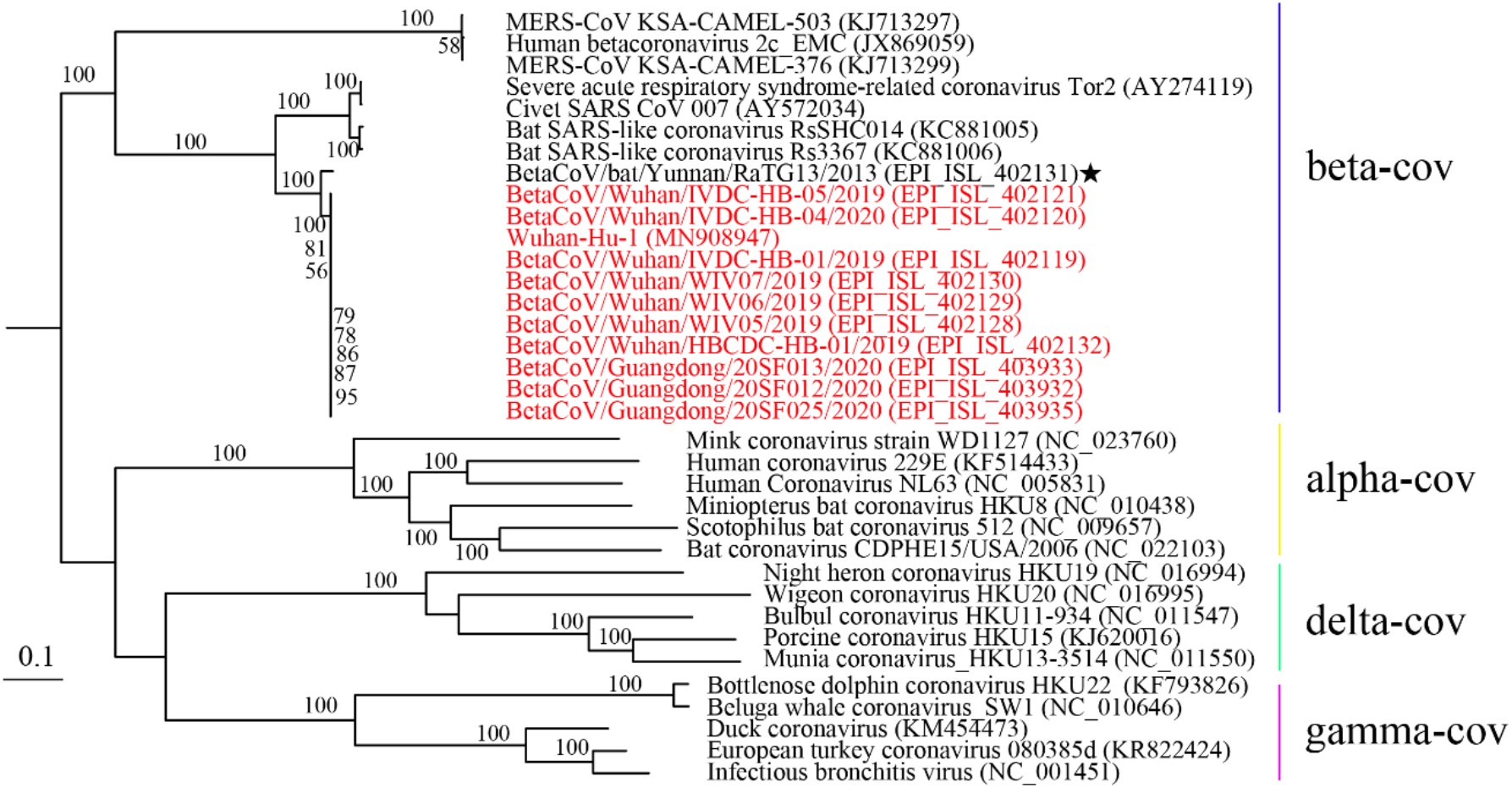
Phylogenetic tree of different CoVs with SARS-CoV-2 based on full genome sequences.

The ML phylogenetic tree demonstrates that CoVs from bat source are found in the inner joint or neighboring clade of SARS-CoV-2. This indicates that bats CoVs particularly Bat/Yunnan/RaTG13 are the source of SARS-CoV-2, and they are emerged and transmitted from bats to humans through some recombination and transformation events in intermediate host.

### Detection of putative recombination within the spike protein

To explore the emergence of SARS-CoV-2 in humans, we investigated CoVs S-protein and its RBD as they are responsible for determining the host range (**Table 1**). The S-protein amino acid sequence identity between SARS-CoV-2 and related beta-CoVs showed that bat/Yunnan/RaTG13 shares highest similarity of 97.43%. However, the amino acid sequence identity of RBD of SARS-CoV-2 with bat/Yunnan/RaTG13 is 89.57%. On the other hand, Beta-CoVs from pangolin sources (pangolin/Guandong/1/2019 and pangolin/Guangdong/lung08) revealed highest RBD amino acid sequence identity of 96.68% and 96.08% respectively with SARS-CoV-2. These indication shows the existence of homologous recombination events within the S-protein gene between bat and pangolin CoVs. Similarity plot analysis of CoVs genome sequences from bat, pangolin and human also indicated a possible recombination within S-protein of SARS-CoVs-19 (**Figure S1**).

**Table 1.** Amino acid sequence identity of S-Protein and RBD of BetaCoVs

| Virus | S protein (%) | RBD(%) |
| --- | --- | --- |
| BetaCoV/pangolin/Guandong/1/2019_EPI_ISL_410721 | 90.02 | 96.68 |
| pangolin/Guangdong/lung08 | 96.3 | 96.08 |
| BetaCoV/bat/Yunnan/RaTG13/2013 _EPI_ISL_402131 | 97.43 | 89.57 |
| BetaCoV/pangolin/Guangxi/P4L/2017_EPI_ISL_410538 | 92.18 | 87.14 |
| BetaCoV/pangolin/Guangxi/P5L/2017_EPI_ISL_410540 | 92.26 | 87.14 |
| BetaCoV/pangolin/Guangxi/P1E/2017_EPI_ISL_410539 | 92.1 | 86.67 |
| BetaCoV/pangolin/Guangxi/P5E/2017_EPI_ISL_410541 | 92.18 | 86.67 |
| BetaCoV/pangolin/Guangxi/P2V/2017_EPI_ISL_410542 | 92.03 | 86.67 |
| SARS_AAR07630.1 | 76.04 | 73.33 |
| Bat_SARS-like_coronavirus_MG772933 | 80.69 | 66.67 |
| MERS_AID55087.1 | 35 | 24.24 |

The amino acid residues change in S-protein of SARS-CoV-2 was further analyzed with SARS-CoV, pangolin and bat CoVs including pangolin/Guandong/1/2019, pangolin/Guangdong/lung08, and bat/Yunnan/RaTG13 (**Figure 2**). Regardless of low homology between SARS-CoV-2 (Wuhan-Hu-1_MN9O8947) and SARS-CoV (SARS_AARO763O), they had many homologues areas in S-protein. The five key amino acid residues of S-protein at positions 442,472, 479,480, and 487 of SARS-CoVs are described to be at the angiotensin-converting enzyme-2 (ACE2) receptor complex interface and supposed to be crucial for human to human and cross-species transmission (Li *et al.,* 2005b; Wu *et al.,* 2012). **Figure 2b** and **Table S1** describe that all key amino acid residues of RBD (except two positions) are completely homologues between SARS-CoV-2 (Wuhan-Hu-1_MN9O8947) and pangolin CoVs (pangolin/Guandong/1/2019 and pangolin/Guangdong/lung08), supporting our postulation of recombination event in S-protein gene. Even though, all five crucial amino acid residues of SARS-CoV-2 for binding to ACE2 are different from SARS-CoV, their hydrophobicity and polarity are similar, having same S-protein structural confirmation and identical RBD 3-D structure (Xu *et al.,* 2020). In addition, six critical key residues in MERS-CoV RBD binding to its receptor dipeptidyl peptidase 4 (DPP4) are all different in SARS-CoV and SARS-CoV-2 related coronavirus **(Figure 2a)**.

**Figure 2.**
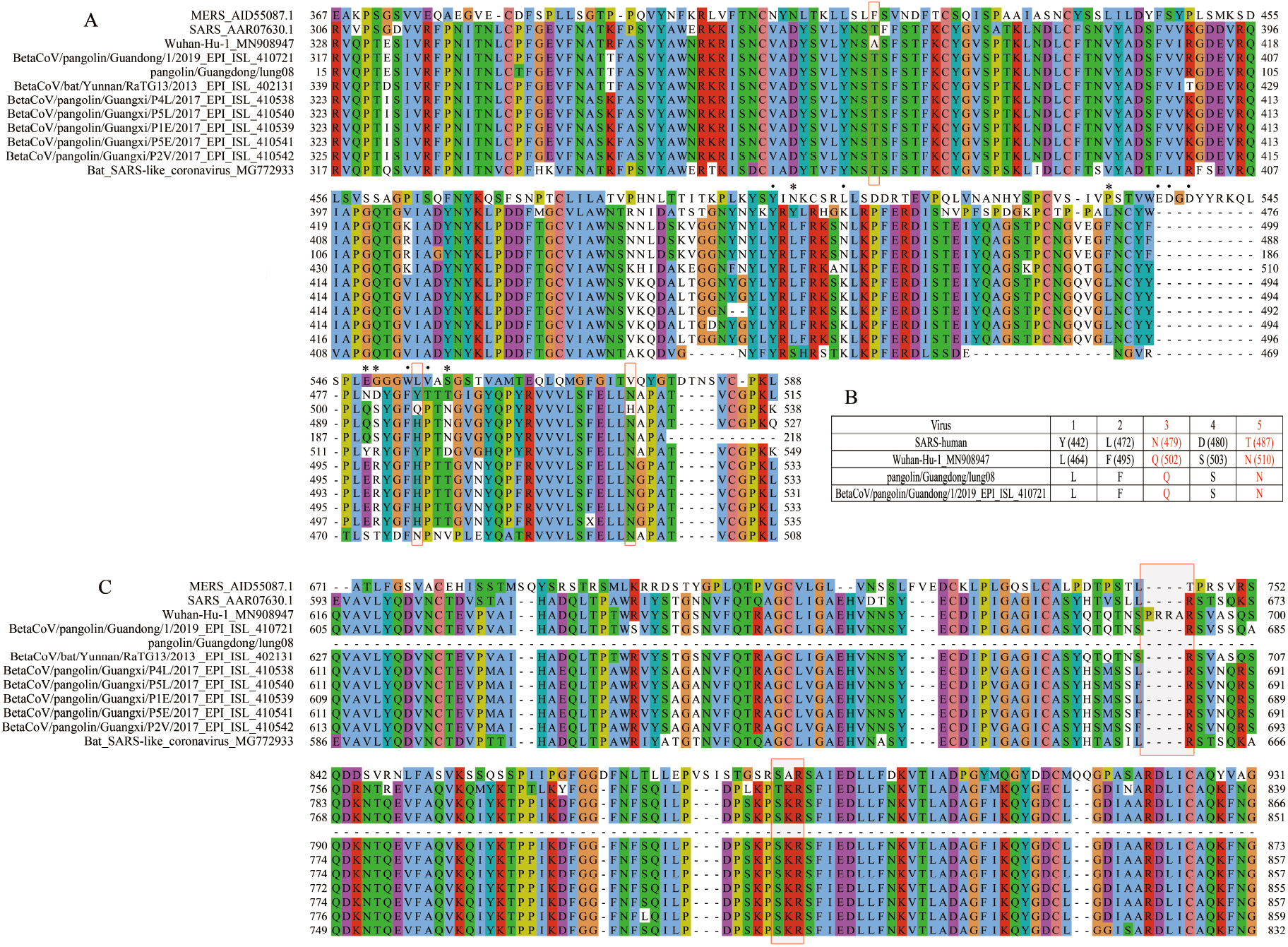
The amino acid residues change in S-protein of CoVs. (A) Amino acid residues change in S1 domain (Asterisks indicate five key amino acid residues of SARS-CoV critical for RBD binding to ACE2; Dots specify critical amino acid residues of MERS-CoV RBD for binding to DPP4). Rectangles specify positions unique in SARS-CoV-2. (B) The combination of five key amino acid residues in SARS-CoVs, SARS-CoV-2, and pangolin CoVs. (C) The amino acid residues change in S2 domain (Rectangles designate changes near two cleavage sites).

### Ongoing mutations in SARS-CoV-2 during its spread

We also investigated some of the important evolutionary and phylogenetic aspects of SARS-CoV-2 during its spread in human population. Mutation in encoding segments of 125 SARS-CoV-2 genomes obtained from public-domain databases were investigated. In comparison with the first reported SARS-CoV-2 (Wuhan-Hu-1_MN9O8947), amino acid substitutions were observed at 87 positions of SARS-CoV-2 open reding frames (orfs) (**Figure 3**). Total number of amino acid substitution in corresponding orfs are listed in **Table 2**. Among different orfs of SARS-CoV-2, orf1a was most variable segment with total number of 44 dissimilar amino acid substitutions. It was followed by spike segment S orf with 13 amino acid residue substitutions. However, orf6 and orf7b are the most conserved regions without amino acid changes. In addition, orf10, E, M and orf7a have tended to be more conserved, with only one or two amino acid substitutions.

**Figure 3.**
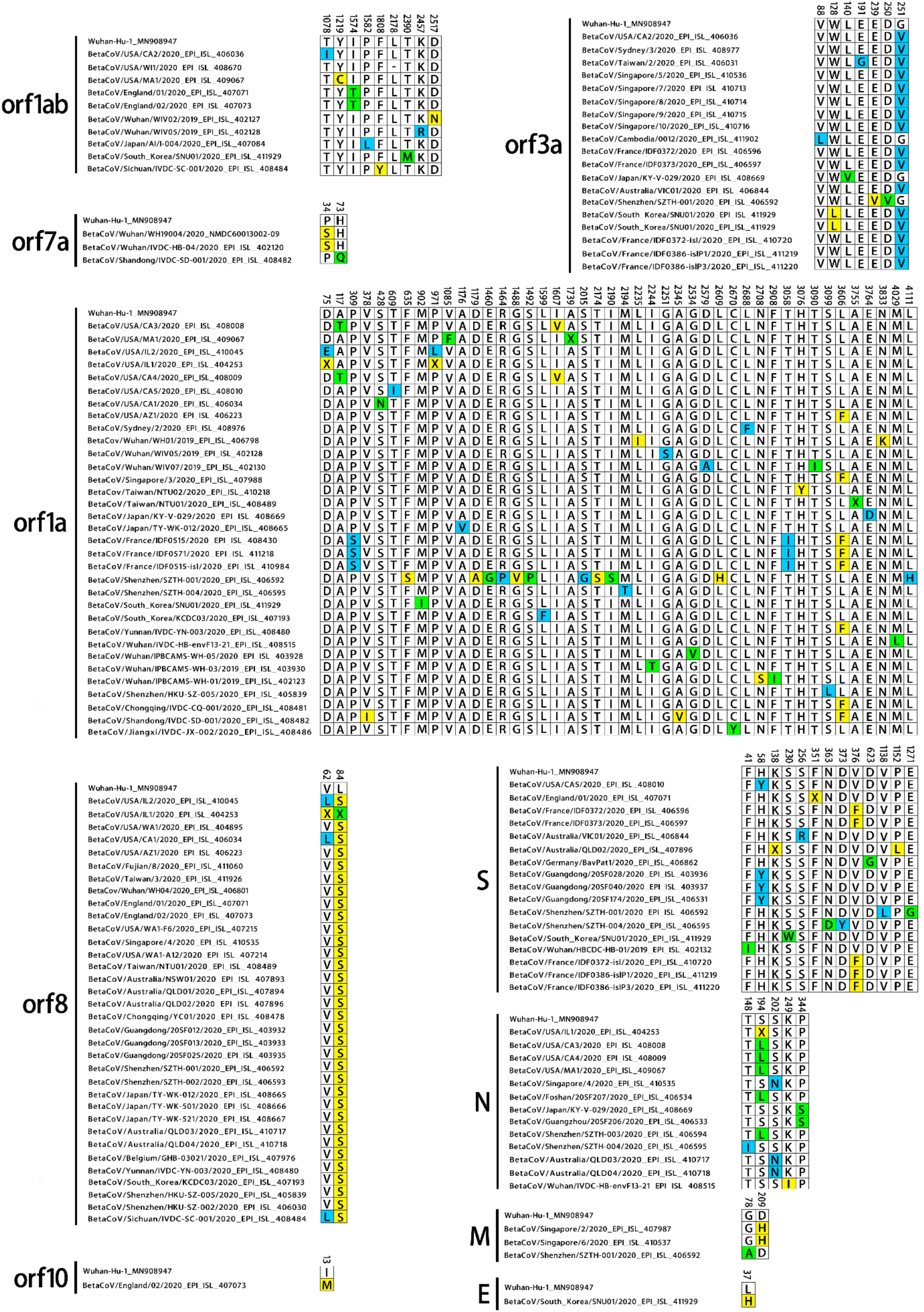
Amino acid substitutions in different open reading frames (orfs) of SARS-CoV-2 (125 genomic sequences collected till February 22, 2020).

**Table 2.** Open reading frame (orf) variation between different SARS-CoV-2

|  | S | E | M | N | orf10 | orf1a | orf1ab | orf3a | orf7a | orf8 |
| --- | --- | --- | --- | --- | --- | --- | --- | --- | --- | --- |
| **Number of dissimilar amino acid substitution** | 13 | 1 | 2 | 5 | 1 | 44 | 8 | 7 | 2 | 4 |

With the global spread of SARS-CoV-2, its amino acid sequence is also significantly varied **(Figure 3).** Usually, RNA viruses have high rate of genetic mutations, which leads to evolution and provide them with increased adaptability (Lin *et al.,* 2019). To further explore SARS-CoV-2 evolution in human, we have performed phylogenetic analysis based on the aforementioned SARS-CoV-2 in correspondence with their amino acid substitution. **Figure 4.** illustrates the phylogenetic tree of SARS-CoV-2 and associated amino acid changes. Our phylogenetic tree demonstrates that BetaCoV/Chongqing/YC01/2020 is the closest SARS-CoV-2 to bat CoV (Bat/Yunnan/RaTG13) as compared to the first reported SARS-CoV-2 (Wuhan-Hu-1_MN9O8947). Taking BetaCoV/Chongqing/YC01/2020 as a reference group, South Korea SARS-CoV-2 (BetaCoV/South_Korea/SNU01/2020) has 7 amino acid substitutions in six orfs (S, E, orf1a, orf31, and orf8). BetaCoV/Shenzhen/SZTH-001/2020 has total of 16 amino acid substitutions in four coding regions with highest substitution of 11 amino acid residues in orf1a. United States of America (USA) SARS-CoV-2 (BetaCoV/USA/MA1/2020) has 5 amino acid substitutions in N, orf1a, orf1ab, and orf8. France CoV (BetaCoV/France/IDF0571/2020) has 3 amino acid mutations in orf1a and 1 amino acid mutation in orf8.

**Figure 4.**
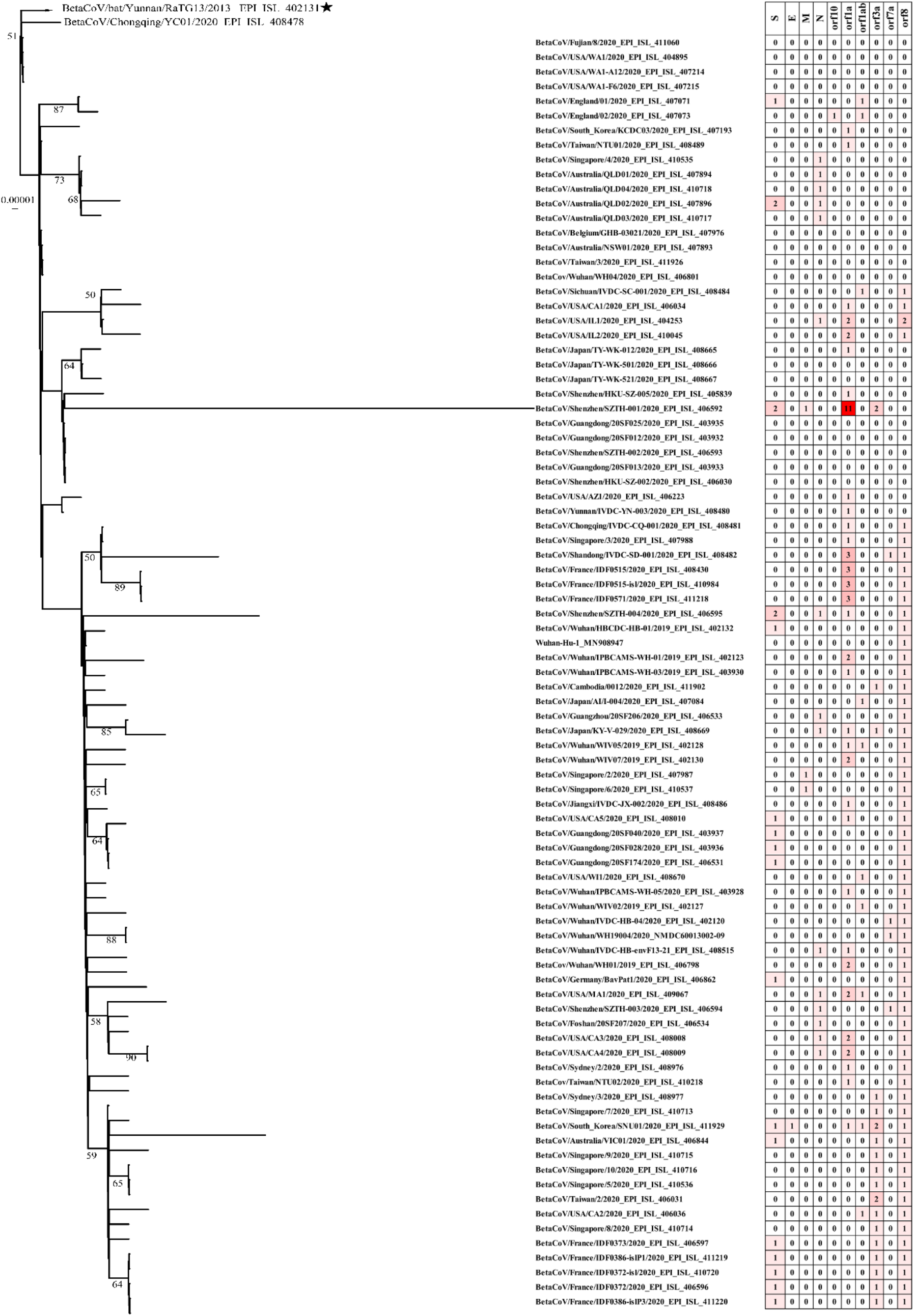
Phylogenetic analysis of SARS-CoV-2 (left) and the number of amino acid substitutions in orfs of corresponding SARS-CoV-2 (right). Bat/Yunnan/RaTG13 was treated as the outgroup.

## CONCLUSION

Based on amino acid and genome sequences analysis and comparison, our results suggest that SARS-CoV-2 is a recombinant virus between bat and pangolin coronaviruses, and the recombination event has been occurred in spike protein genes. Our finding suggest that pangolin is the most possible intermediate SARS-CoV-2 reservoir, which may have given rise to cross-species transmission to humans. These new findings suggest further research to investigate pangolin as a SARS-CoV-2 reservoir. Another important outcome of our analysis is the genetic mutations and evolution of SARS-CoV-2 as it spread globally. These findings are very significant for controlling the SARS-CoV-2 pandemic.

## ACKNOWLEDGMENT

This work was supported by the National Natural Science Foundation of China grants 31770948 and 31570875 and Marine Economic Development Special Fund of Fujian Province (FJHJF-L-2020-2) and the High-level personnel introduction grant of Fujian Normal University (Z0210509). We also gratefully acknowledge the support from Key Research and Development Program (COVID-19) of Hainan (No. ZDYF(XGFY)2020002).

## MATERIALS AND METHODS

### Sequence data collection

One hundred and twenty-five newly sequenced SARS-CoV-2 complete genomes were obtained from Global Initiative on Sharing All Influenza Data EpiFluTM database (GISAID EpiFlu^™^) and GenBank. Closely related beta-CoVs genomes sequences from different hosts were also collected and analyzed together with SARS-CoV-2. Open reading frames (orfs) of CoVs genomes were predicted using ORFfìnder (v0.4.3) with default parameters ignoring nested orfs.

Raw pair-end reads of pangolin dataset sample (SRR10168377) obtained from NCBI were filtered with bbmap.sh (v38.79) by removing adaptors, trimming low quality reads from both sides (quality value < 20), and reads length less than 50 nt were ignored. Host reference genome (pangolin ManJav1.0, GCF_001685135.1) contaminant reads were removed by bowtie2 (v2.3.5.1) [13]. Pangolin CoV genome fragments were assembled via MEGAHIT (v1.2.9) (Li *et al*., 2015).

### Phylogenetic and recombination analysis

The sequences of CoVs were aligned using multiple sequence alignment MAFFT (v7.450) (Katoh et al., 2019). Aligned sequences were visualized with Jalview (v2.10.3) (Waterhouse *et al.,* 2009). Poorly aligned regions and gaps were removed by trimAL (v1.4.rev22) (Capellagutiérrez *et al*., 2009). Maximum likelihood (ML) phylogenetic trees of whole genome sequences were constructed in IQ-TREE (v1.6.12) (Nguyen *et al*., 2015). Support for inferred relationships in the phylogenetic tree was assessed by bootstrap analysis with 1000 replicates and the best-fit substitution model was determined by IQ-TREE model test.

## SUPPLEMENTAL INFORMATION

**Figure S1.**
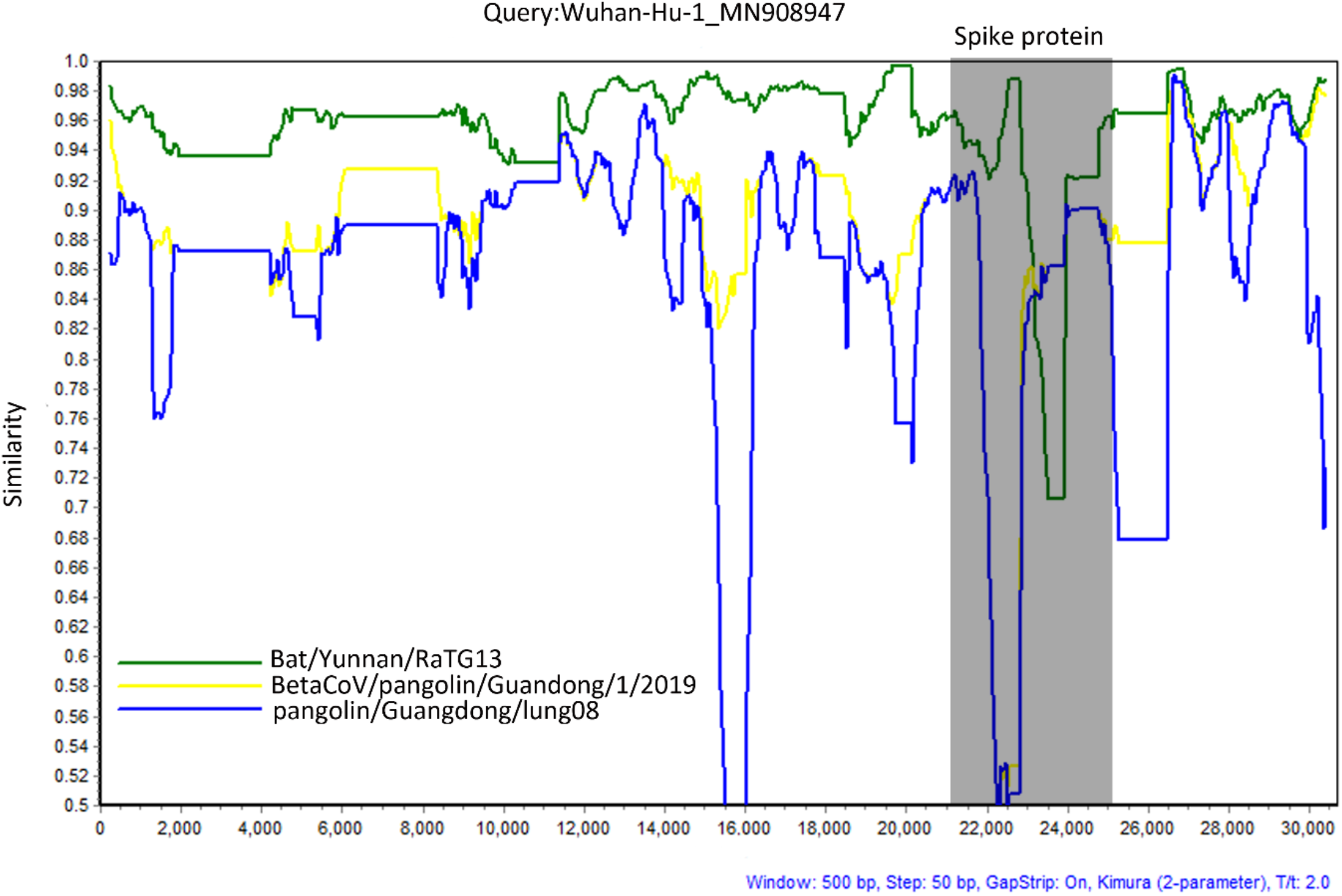
Similarity plot of the full genome sequence among Bat/Yunnan/RaTG13, BetaCoV/pangolin/Guandong/1/2019, pangolin/Guangdong/lung08 and Wuhan-Hu-1_MN9O8947 (as the query). Grey box highlights the Spike protein coding region, supposed to be involved in recombination. Similarity scores between genomic sequences were generated by Simplot (v3.5.1) (Lole *et al.,* 1999).

**Table S1.** The interactions of SARS-CoV RBD with ACE2 and their comparison with RBD of SARS-Cov-2, BetaCoV/pangolin/Guandong/1/2019 and pangolin/Guangdong/lung08.

|  | SARS-CoV | ACE2 | SARS-Cov-2 | BetaCoV/pangolin/Guandong/1/2019 | pangolin/Guangdong/lung08 |
| --- | --- | --- | --- | --- | --- |
| Hydrogen bonds | N473(ND2) | Q24(OE1) | N496 | N485 | N183 |
|  |  | E35(OE1) | Q502 | Q491 | Q189 |
|  | Y491(OH) | E37(OE1) | Y514 | Y503 | Y201 |
|  |  | E37(OE2) | Y514 | Y503 | Y201 |
|  | Y436(OH) | D38(OD1) | Y458 | Y447 | Y245 |
|  | Y436(OH) | D38(OD2) | Y458 | Y447 | Y245 |
|  | T486(OG1) | Y41(OH) | T509 | T498 | T296 |
|  | T487(N) | Y41(OH) | N510 | N499 | N297 |
|  |  | Q42(NE2) | Q507 | H496 | H294 |
|  |  | Q42(NE2) | G455 | G444 | G242 |
|  |  | Q42(NE2) | Y458 | Y447 | Y245 |
|  | Y436(OH) | Q42(OE1) |  |  |  |
|  | N473(ND2) | Y83(OH) | N496 | N485 | N183 |
|  | Y475(OH) | Y83(OH) | Y498 | Y487 | Y185 |
|  |  | Q325(OE1) |  |  |  |
|  |  | R426(NH2) |  |  |  |
|  |  | N330(ND2) |  |  |  |
|  | T486(O) | K353(NZ) | Y504 | Y493 | Y291 |
|  |  | K353(NZ) | G505 | G494 | G292 |
|  | G488(N) | K353(O) | G511 | G500 | G198 |
|  |  | R393(NH2) | Y514 | Y503 | Y201 |
| Salt bridges |  | D30(OD2) | K426 | R415 | R113 |
|  | R426(NH1) | E329(OE2) |  |  |  |
|  | R426(NH2) | E329(OE2) |  |  |  |

Red color highlights interaction positions of SARS-CoV-2 and pangolin CoVs with different amino acids residues.

